# Characterizing human CMV-specific CD8^+^ T cells using multi-layer single-cell omics

**DOI:** 10.1101/2024.09.17.613216

**Authors:** Ioanna Gemünd, Lorenzo Bonaguro, Matthias Becker, Sophie Müller, Clemens Joos, Elena De Domenico, Anna C Aschenbrenner, Joachim L Schultze, Andreas Moosmann, Marc D Beyer

## Abstract

In this study we established a comprehensive workflow to collect multi-omics single-cell data using a commercially available micro-well based platform. This included whole transcriptome, cell surface markers (targeted sequencing-based cell surface proteomics), T cell specificities, adaptive immune receptor repertoire (AIRR) profiles and sample multiplexing. With this technique we identified novel paired T cell receptor sequences for three prominent human CMV epitopes. In addition, we review the ability of dCODE dextramers to detect antigen-specific T cells at low frequencies by estimating sensitivities and specificities when used as reagents for single-cell multi-omics.

**Motivation:** In this study, we report the first five-layer multi-omics dataset using the BD Rhapsody single-cell platform for the characterization of human antigen-specific T cells. Modalities include whole transcriptome, T cell receptor (TCR) sequences, T cell antigen specificity measured by dCODE dextramers, surface marker proteins and combinatorial sample multiplexing combining two distinct hashing approaches.

## 4. Introduction

Improving our understanding of immune responses requires sophisticated methods capable of capturing multiple layers of cellular information simultaneously. The emergence of advanced single-cell multi-omics technologies has revolutionized our understanding of cellular heterogeneity and immunological processes ^1,2^. Recently, new techniques have enabled the in-depth characterization of lymphocytes and their diverse adaptive immune receptor repertoires (AIRR). This includes profiling of immune receptor sequences at the transcript level as well as antigen interaction on the protein level using DNA-barcoded peptide-MHC (pMHC) or antigen multimers for T cells or B cells, respectively ^3^. Combined with other single-cell modalities like transcriptomics or targeted sequencing-based proteomics, AIRR information allows for the characterization of intra- and interclonal phenotypic heterogeneity of lymphocytes in the context of microenvironmental cues as well as clonal tracking of cells over time and across tissues ^4–6^. This is important as T cell clones can exhibit varied functionalities and phenotypes based on their peptide specificities, MHC restriction, tissue localization, and antigen exposure ^7–9^.

In this work, we present an exemplary workflow for collecting paired single-cell data encompassing whole transcriptome, 31 surface markers, immune receptor sequences, antigen specificities, and combinatorial multiplexing using the BD Rhapsody single-cell platform ^10^. We utilize this cutting-edge technology to identify and characterize novel T cell receptor (TCR) sequences of human cytomegalovirus (CMV)-specific CD8^+^ T cells. Additionally, we address technical aspects crucial for planning multi-omics experiments that include AIRR information.

When dealing with biological materials containing T cells, the identities and proportions of T cells specific for a certain antigen within a sample are hard to estimate. Factors such as the organ from which a sample was derived as well as disease status and age of the donor add additional variability. In many cases, frequencies of antigen-specific T cells of interest will be low. To study these cells, enrichment methods like fluorescence-activated cell sorting (FACS) with pMHC multimers are often required, as they improve the detection of low-abundance transcripts and proteins in rare antigen-specific cell populations and allow sequencing experiments to be conducted in a cost-effective manner. However, additional processing steps can be inconvenient or unfeasible in some experimental setups and may even alter cellular states. More importantly though, prior pMHC multimer enrichment might not fully capture the entire T cell response to a given antigen ^11^. This limitation can result in the failure to notice responses to uncharacterized MHC-presented peptide epitopes and missing out on analyzing T cells in the context of their overall immune landscape. Therefore, in studies involving complex immunophenotypes or exploratory research, barcoded pMHC multimer staining - either without subsequent flow cytometry enrichment or with simultaneous analysis of both enriched and unenriched cells - can enable comprehensive, side-by-side characterization of gene expression and TCR profiles for antigen-specific T cells with both known and, with the advancement of TCR-antigen specificity inference tools, unknown specificities for which potential reactivity can be computationally predicted ^12,13^.

This raises the question whether DNA-barcoded pMHC complexes, such as dCODE dextramers, can reliably identify antigen-specific T cells at low frequencies for single-cell transcriptomics studies. In this study we address this through a spike-in experiment using human CMV-specific T cells with known peptide-MHC specificities and compare the performance of dCODE dextramers when used as sequencing reagents against conventional flow cytometry reagents. Taken together, the presented workflow offers significant potential for studying antigen-specific T cells in various biological contexts and provides a valuable resource for researchers using the BD Rhapsody single-cell platform or considering experiments with dCODE dextramers.

## 5. Results

### Single-cell multi-omics analysis allows for the delineation of antigen-specific T cells and background samples

T cell immunity is characterized by diverse functionalities and phenotypes among different T cell clones and single-cell multi-omics approaches enable comprehensive exploration of this diversity. Here we give a framework of how this can be technically achieved employing a five-layer BD Rhapsody multi-omics approach that includes whole transcriptome analysis (WTA), T cell receptor (TCR) sequencing, antibody sequencing (AbSeq), dCODE dextramers, and dual hierarchical sample multiplexing (Fig. 1A and Supp. Fig. 1).

**Figure 1.**
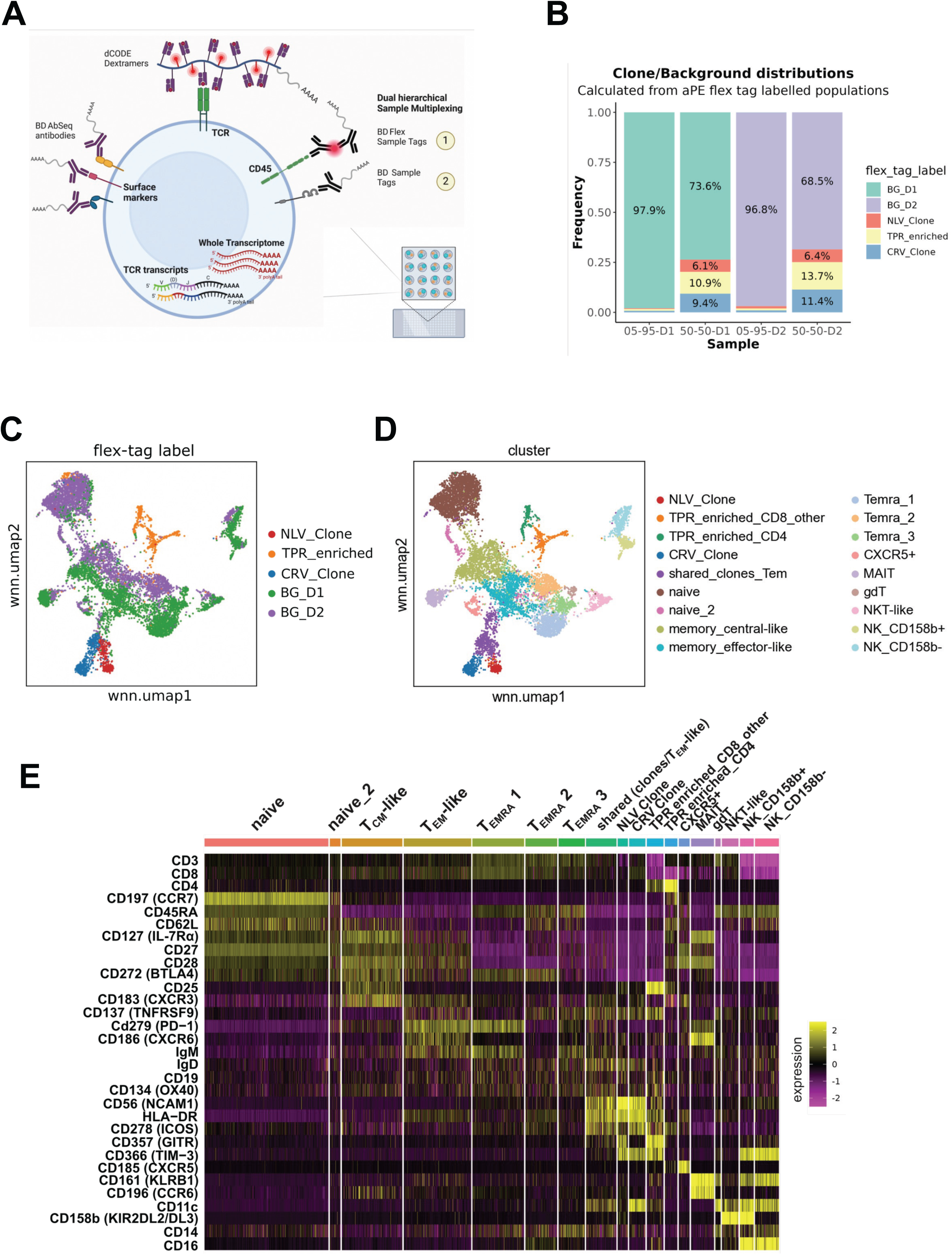
Single-cell multi-omics workflow allows delineation of spike-in and background samples. (A) Scheme of different multi-omics modalities that are captured per cell using the BD Rhapsody single-cell platform. (B) Distribution of spike-in and background cells in the four dCODE dextramer staining conditions. (C-D) wnnUMAP representation of single-cell data with coloring according to flex sample tag label (origin of five input populations) and cell clustering. (E) Expression heatmap of 31 surface markers by cluster (coloured by z-transformed expression).

As a proof-of-concept to determine the ability to identify cytomegalovirus (CMV)-specific T cells at low frequency using dCODE dextramers, we conducted a spike-in experiment. As spike-in, a mixture of three samples enriched for CMV-specific CD8^+^ T cells in a ratio of 1:1:1 was used. This included two previously described T cell clones ^8,14^ and one sample of PBMCs 10 days after peptide stimulation. In the following these samples will be called ‘NLV clone’, ‘CRV clone’ and ‘TPR enriched’ according to the CMV peptides they recognize: NLVPMVATV (pp65_495-503_) for the NLV clone, CRVLCCYVL (IE-1_309-317_) for the CRV clone, and TPRVTGGGAM (pp65_417-426_) for the TPR enriched sample, which was derived from PBMCs of a healthy donor (HLA-B*07:02^+^). Correspondingly, we utilized three dCODE dextramers targeting the respective CMV epitopes: DexA (pp65_495-503_ on HLA-A*02:01), DexB (pp65_417-426_ on HLA-B*07:02), and DexC (IE-1_309-317_ on HLA-C*07:02). A negative control dextramer (DexN) carrying a nonsense peptide was also employed for later background correction (HLA-B*08:01/AAKGRGAAL). Prior to the spike-in experiment, individual flow cytometry stainings with dCODE dextramers (PE labeled) revealed that the NLV clone binds to DexA, with 97% of cells being DexA-PE positive. Similarly, 98% of the CRV clone cells were DexC-PE positive (Sup. Fig. 2A, B). The TPR enriched sample contained about 16% DexB-PE^+^ cells and approximately 18% non-T cells (Sup. Fig. 2B), reflecting that this sample originated from PBMCs 10 days after peptide stimulation.

The three-sample spike-in mix was compared against polyclonal CD8^+^ T cells from PBMCs of two healthy donors, in the following referred to as ‘background’. Specifically, DexA was HLA-matched with background donor 1 (BG_D1) (HLA-A02:01^+^; HLA-B07:02^-^ and HLA-C*07:02^-^), while DexB and DexC were HLA-matched with background donor 2 (BG_D2) (HLA-B07:02^+^ and HLA- C*07:02^+^; HLA-A02:01^-^). To estimate the sensitivity and specificity of the capture of antigen-specific T cells with dCODE dextramers in our multi-omics approach, we needed a low-frequency spike-in condition. For this we chose to perform a 5% spike-in of the aforementioned three-sample spike-in mix, meaning that each sample’s actual spike-in frequency was about 1.67% (Table 1). Additionally, we included a 50% spike-in plus 50% background condition to ensure sufficient cells for characterization of the spike-in mix (Supp. Fig. 1, Table 1).

**Table 1.** Spike-in and multiplexing strategy for CMV multi-ome experiment. Overview of the used indexing strategy. A hierarchical cell labelling system (first stained with flex sample tags, then stained with standard sample tags) allows to allocate each cell to both the corresponding input material (BG_D1, BG_D2, NLV Clone, TPR enriched and CRV clone), as well as to one of the four staining mixes (50-50-D1, 5-95-D1, 50-50-D2 and 5-95-D2). Flex and standard sample tags use different DNA barcodes.

| Sample | BG/<br>Spike-In | Percentage of cell in<br>sample | Cells | First cell label<br>(Flex sample tag) | Second cell label<br>(Standard sample tag) |
| --- | --- | --- | --- | --- | --- |
| 1 | 95 % | 95 % | BG_D1 | 1 | 1 |
|  | 5 % | 1.67 % | NLV | 3 | 1 |
|  |  | 1.67 % | TPR | 4 | 1 |
|  |  | 1.67 % | CRV | 5 | 1 |
| 2 | 50 % | 50 % | BG_D1 | 1 | 2 |
|  | 50 % | 16.7 % | NLV | 3 | 2 |
|  |  | 16.7 % | TPR | 4 | 2 |
|  |  | 16.7 % | CRV | 5 | 2 |
| 3 | 95 % | 95 % | BG_D2 | 2 | 3 |
|  | 5 % | 1.67 % | NLV | 3 | 3 |
|  |  | 1.67 % | TPR | 4 | 3 |
|  |  | 1.67 % | CRV | 5 | 3 |
| 4 | 50 % | 50 % | BG_D2 | 2 | 4 |
|  | 50 % | 16.7 % | NLV | 3 | 4 |
|  |  | 16.7 % | TPR | 4 | 4 |
|  |  | 16.7 % | CRV | 5 | 4 |

In order to later reidentify all five input populations (three CMV-enriched populations and two background donors), as well as four dCODE dextramer staining conditions (5 % and 50 % CMV spike-in mix against either BG_D1 or BG_D2), we employed a dual hierarchical indexing strategy. Individual input populations were labeled with anti-PE BD flex tags binding PE-labeled anti-CD45 in the primary staining (Sup. Fig. 1, Part A). Subsequently, the four spike-in/donor mixes were labeled with regular BD sample tags (Sup. Fig. 1, Part B). This allowed for the simultaneous allocation of spike-in input samples and staining mixes and facilitated the estimation of dextramer performance within each setting.

We loaded 20,000 cells on two BD cartridges (technical replicates), performed the BD Rhapsody workflow and proceeded to bioinformatic analysis after sequencing and preprocessing (see methods and Figure S5 for details). The first step of the analysis was assigning cells to their corresponding input material (flex sample tag for BG_D1, BG_D2, NLV_Clone, TPR_enriched, or CRV_Clone) and staining mix (standard sample tag for 5:95 or 50:50 of spike-in/background with either background donor 1 or 2) (Table 1). Cells were retained for analysis only if both flex tag and sample tag labels, thus information on both input material and staining mix, were present, resulting in a dataset of 9,771 cells. After demultiplexing, the distribution of the five input samples within staining mixes showed a ratio of approximately 3%/30% spike-in and 97%/70% background cells per staining condition (Figure 1B), indicating lower than expected spike-in frequencies likely due to lower viability of the clones after cell thawing.

We utilized uniform manifold approximation and projection (UMAP) to reduce data dimensionality^15^. A weighted nearest neighbor (wnnUMAP) approach was chosen for multimodal UMAP embedding, based on paired scRNA-seq and AbSeq information ^16^. This approach allowed cells of the same subset or type to accumulate in specific UMAP clusters, indicating similar transcriptome and surface marker expression. Cell clustering categorized the dataset into 18 main clusters (Figure 1D), which we further analyzed by identifying cell-specific markers combining transcriptome and surface molecule information (Figure 1E, Sup. Fig. 2D). No technical batch effects from both cartridges were observed (Sup. Fig. 2C). Further analysis revealed that background samples from different donors were distinguishable in the wnnUMAP (Figure 1C). Background CD8^+^ T cells clustered into four main cell states: naïve, central memory-like (T_CM_-like), effector memory-like (T_EM_-like), and effector memory T cells re-expressing CD45RA (T_EMRA_) (Figure 1D). Naïve T cells were characterized by CCR7 gene expression and surface markers CCR7, CD62L, and CD45RA (Figure 1E, Sup. Fig. 2D). T_CM_-like cells expressed CD127, CD27, and CD28, with intermediate levels of CCR7 and CD62L, and no CD45RA (Figure 1E). The T_EM_-like cells exhibited high expression of PD-1 (CD279) and CXCR6, as well as genes encoding for *GZMK* and *GZMA* required for cytotoxic function (Figure 1E, Sup. Fig. 2D) while cells lacked the surface markers CD62L, CCR7, and CD45RA. T_EMRA_ cells re-expressed CD45RA and lacked CD127 and CD27 (Figure 1E). Additional smaller clusters included MAIT cells, γδ T cells, CXCR5^+^ cells, NKT cells, and NK cell clusters (Figure 1D).

The NLV clone, TPR enriched sample, and CRV clone clustered separately from the CD8 polyclonal background samples (Figure 1C). A shared cluster of clone and T_EM_-like cells with high mitochondrial gene expression was observed (“shared_clones_Tem”). The TPR enriched sample appeared more heterogeneous and also included CD4^+^ T cells (Figure 1E). Both NLV and CRV clones, along with the TPR fraction containing CD8^+^ T cells and the shared clone/T_EM_-like cluster, exhibited high surface expression of activation markers HLA-DR, ICOS, and CD56 (NCAM1) (Figure 1E). The NLV and CRV clones were further distinguished by increased gene expression of CD8^+^ T cell activation markers *IFNG*, *SLAMF7*, *GZMA* and *GNLY* (Sup. Fig. 2D). This supported the notion that the antigen-specific CD8^+^ T cell clones were of CD8^+^ T cell origin but transcriptionally different from *ex viv*o isolated CD8^+^ T cells, likely due to their *in vitro* cultivation and possibly also influenced by the distinct activation and differentiation history of CMV-specific CD8^+^ T cells *in vivo*.

### Characterization of CMV-specific CD8^+^ T cells

To identify dextramer-positive (Dex^+^) cells, we analyzed the distribution of normalized expression for each dextramer to establish a reasonable cutoff. The data revealed a clear separation into Dex^high^ and Dex^low^ populations (Sup. Fig. 3A). The cutoff was set to divide these populations at local minima of the density plots, which displayed a multimodal distribution. Thresholds for Dex^+^ cells were automatically defined using k-means clustering (Supp. Fig. 3A).

We then investigated which clusters contained Dex^+^ cells (Figure 2A). As expected, the majority of the NLV clone was defined as DexA^+^, and the majority of the CRV clone as DexC^+^. The shared clone and T_EM_-like cell cluster also included DexA^+^ and DexC^+^ cells. Additionally, a fraction of the ‘TPR_enriched_CD8/other’ cluster was DexB^+^. Background donor 1 (BG_D1), which was HLA-matched to DexA, displayed a distinct fraction of DexA^+^ cells, whereas background donor 2 (BG_D2), HLA-matched to DexB and DexC, showed a small cluster of DexB^+^ and some interspersed DexC^+^ cells (Figure 2B). Dex^+^ cells in the background samples mainly localized to the T_EMRA_ and T_EM_-like clusters (Figure 2B). These cells potentially correspond to endogenous antigen-specific memory CD8^+^ T cells in these donors (Sup. Fig. 3C).

**Figure 2.**
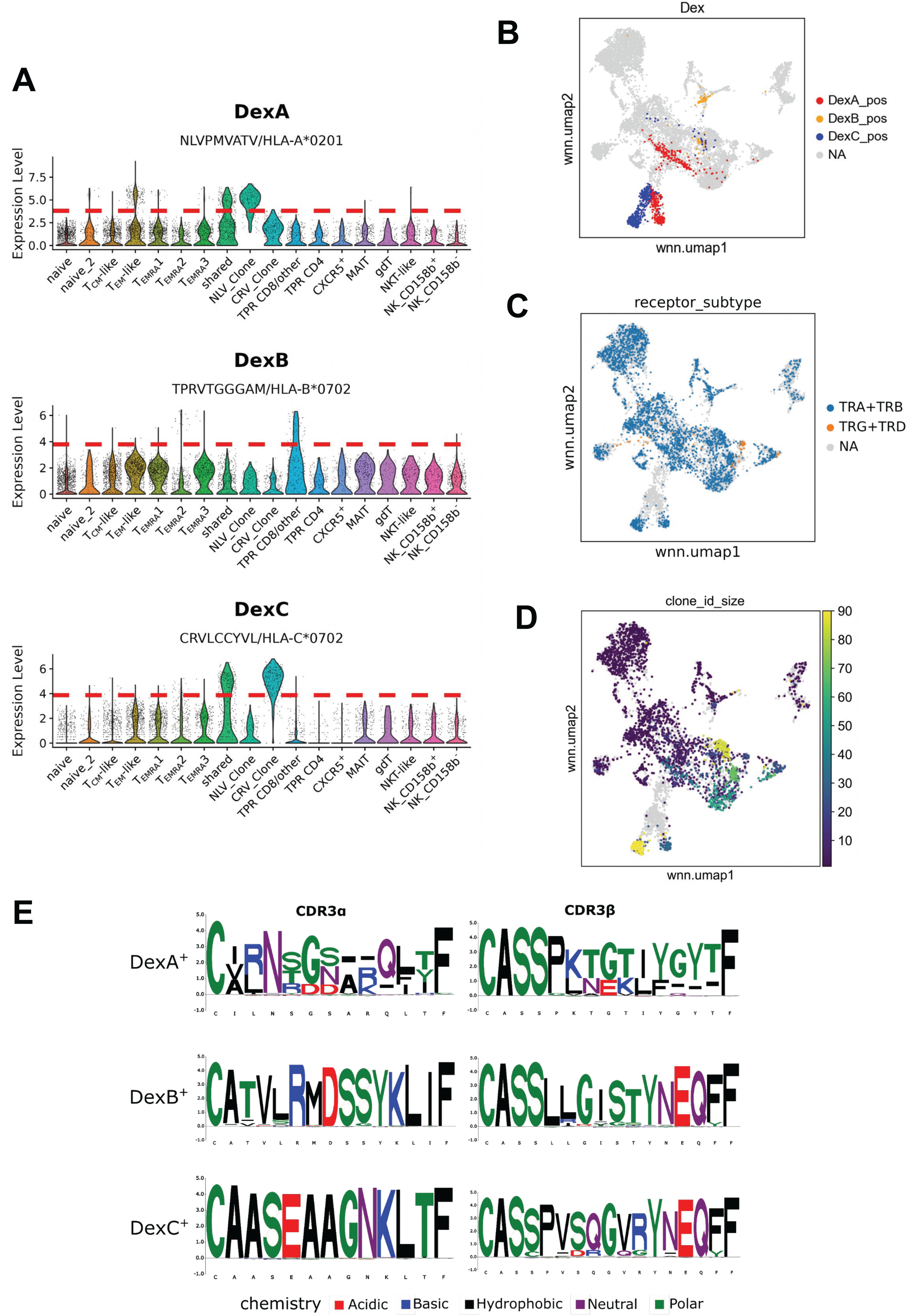
Characterization of CMV-specific CD8^+^ T cells. (A) Distribution of noise-corrected CLR-normalized DexA, DexB and DexC counts per cluster. Red line indicates cutoff for Dex-positive label. (B-D) wnnUMAP representation of dataset with highlighted DexA^+^, DexB^+^ and DexC^+^ cells (B), colored by receptor type after filtering for ‘single pair’, ‘extra VJ’ and ‘extra VDJ’ sequences (C) and showing the corresponding clone size for each cellular TCR sequence (D). (E) CDR3 sequence logo plots depicting under- and over-represented amino acids of TCR CDR3α and CDR3β amino acid sequences. Logo plots are shown for DexA^+^, DexB^+^ and DexC^+^ cells. Colors indicate amino acid chemistry.

Subsequently, we examined the occurrence of CMV-specific CD8^+^ T cells under different dCODE dextramer staining conditions. The frequencies of Dex^+^ cells in the four spike-in/background staining conditions for each dextramer are shown in Supp. Fig. 3B. Samples with HLA-matched backgrounds consistently contained higher frequencies of Dex^+^ cells compared to HLA-mismatched samples. However, even in samples with low-frequency spike-ins, Dex^+^ cells were consistently detectable (Sup. Fig. 2B), indicating that the assay was sensitive enough to detect antigen-specific T cells across all spike-in frequencies observed in our data.

As it has been reported that HLA-C multimers like DexC can bind to CD158b (KIR2DL2/3) on CD8^+^ T cells with low affinity in a CMV-epitope-independent manner ^17^, we also investigated the co-expression of CD158b (KIR2DL2/3) and DexC. An oligo-coupled CD158b antibody was included to identify potential epitope-independent KIR2DL2/3 binding of DexC. Co-expression analysis did not reveal a KIR2DL2/3^+^ DexC^high^ population (Sup. Fig. 3D), indicating that DexC specifically bound to TCRs targeting the IE-1_309-317_ epitope.

To evaluate the performance of dCODE dextramers as oligo-coupled reagents read out by sequencing, we compared their performance to flow cytometry data from the same reagents and conditions. In the flow cytometry data, DexA and DexC molecules bound to nearly the entirety of the corresponding clones (Supp. Fig. 2B). We used this as a reference to estimate the sensitivity and specificity of DexA and DexC in the single-cell RNA-seq protocol. Cells from the NLV or CRV clones stained with flex tags were assumed capable of binding DexA and DexC, respectively, while DexB was excluded due to the heterogeneity of the TPR enriched sample.

We assessed the overlap between DexA^+^ and DexC^+^ cells based on dextramer detection (“predicted class”) and cells labeled as NLV or CRV clones by flex tags (“actual class”) using a confusion matrix, and calculated sensitivity and specificity based on this comparison (Table 2). For DexA, the positive “actual class” included the NLV clone, with the negative class consisting of cells either HLA-mismatched to DexA or cells from the CRV clone. For DexC, the positive “actual class” were cells from the CRV clone, and the negative class contained cells HLA-mismatched to DexC or cells from the NLV clone. The predicted classes were previously defined based on detected CLR-normalized dextramer sequencing counts (Dex^+^ and Dex^-^ labels). Using these definitions, we observed that DexA and DexC exhibited specificities of 100% and 99.9%, respectively, indicating a minimal false-positive rate. Sensitivities were 81.2% for DexA and 83.1% for DexC (Table 2).

**Table 2.** Dextramer performance estimation for DexA and DexC. Sensitivity and specificity for performance of dextramer DexA on the NLV clone sample and DexC on the CRV clone sample when used as sequencing reagents. These values were calculated using a confusion matrix that compared the ‘actual’ class label (as defined by the flex tag) with the ‘predicted’ class (Dex^+^/Dex**^-^** label). Dex^+^ cells originating from a clone sample are true positives (TP), while Dex**^-^**labelled cells of the same origin are false negatives (FN). Dex**^-^**cells originating from the opposite clone or HLA-mismatched background sample are true negatives (TN), while Dex^+^ cells of this origin are false positives (FP).

|  | DexA & NLV clone | DexC & CRV clone |
| --- | --- | --- |
| Sensitivity | 0.812 | 0.831 |
| Specificity | 1.000 | 0.999 |

Taken together, this demonstrates that dCODE dextramers can be highly specific and reasonably sensitive when used as oligo-coupled reagents for single-cell RNA-seq, providing reliable identification of antigen-specific T cells. However, differences in sensitivity between single-cell RNA-seq and flow cytometry exist, highlighting the need for careful consideration when interpreting results across modalities.

### TCR repertoires of CMV-specific CD8^+^ T cells

Analysis of the single-cell immune receptor data identified 7,499 cells with both immune receptor sequences and corresponding transcriptome data. Filtering the TCR dataset revealed that 7,281 of these cells had TCR sequences, while the remainder were BCR-expressing (2 cells) or multi-chain cells expressing more than two TCRα- or two TCRβ-chains (213 cells). For approximately 25% of cells in the transcriptome data, single pairs of TCRα/β or TCRγ/δ chains could be reconstructed (Supp. Fig. 4A, Fig. 2C). For the majority of cells at least one TCRα or TCRβ chain was detected (6,561 in total), while the remaining cells either had TCRγ/δ chain (270 cells) or ambiguous TCR combinations, such as TCRβ+γ (450 cells). Notably, TCRγδ sequences co-localized with the γδ T cell cluster identified in the scRNA-seq dataset (based on *TRGC2* & *TRDC* gene expression) (Fig. 2C, Supp. Fig. 2D). Interestingly, the shared cluster between clones and T_EM_-like cells contained very few cells with TCR sequences (Fig. 2C). Clone sizes, defined by identical nucleotide TCR sequences, were highest in the T_EMRA_ populations and the NLV and CRV clones (Fig. 2D). The clone size of the CRV clone appeared larger than of the NLV clone because the latter consisted of a population with a shared TCRβ chain but two different TCRα chains (Table 3, clonotype cluster 1 and 15; Suppl. Fig. 4B).

**Table 3.**
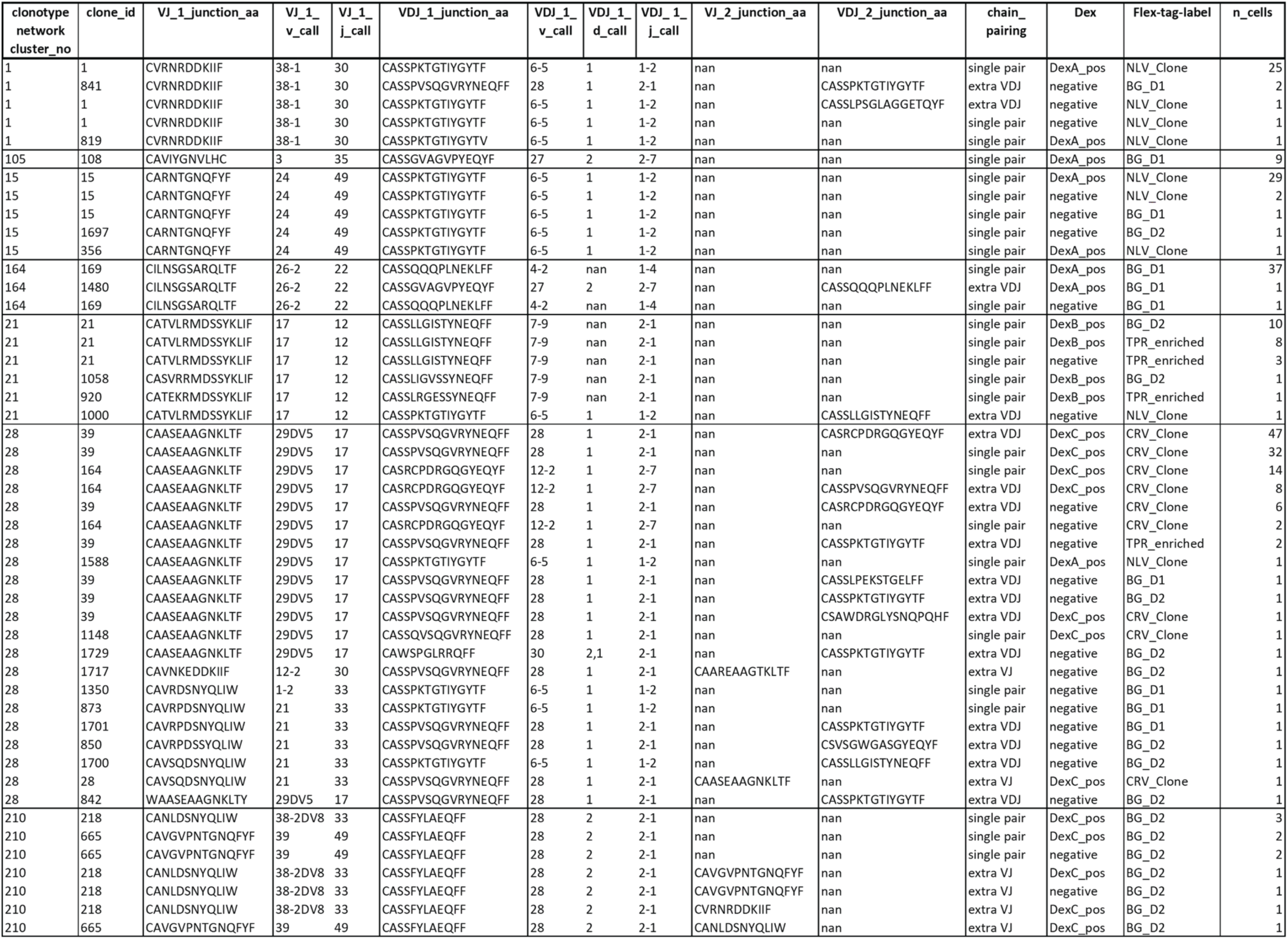
CDR3 amino acid sequences and TCR gene usage in selected clonotype clusters. The table lists CDR3 amino acid sequences, TCR gene usage, clone frequency, clone_id, dextramer binding (“Dex”), and flex tag labels for clones within selected clonotype clusters (cluster numbers are indicated in the first column). The “Dex” column denotes whether a T cell bound to a specific dextramer, indicating antigen specificity for the peptide-MHC (pMHC) complex presented by that dextramer (DexA: HLA-A*02:01/NLVPMVATV(pp65_495-503_); DexB: HLA-B*07:02/TPRVTGGGAM (pp65_417-426_); DexC: HLA-C*07:02/CRVLCCYVL (IE-1_309-317_). The ‘clone_id’ label refers to cells that share identical TCR nucleotide sequences, while the ‘clonotype_cluster’ label indicates groups of clones that share a high degree of CDR3 amino acid similarity between their VJ and VDJ chains.

Next, we generated CDR3 amino acid logo plots for flex tag labels (NLV, CRV, TPR; Suppl. Fig. 4B) versus Dex-positive cells (DexA, DexC, DexB; Fig. 2E). This revealed a nearly identical CDR3 amino acid sequence for the CRV clone (DexC), while CDR3 amino acid sequences of the NLV clone and the TPR enriched sample differed from DexA^+^ and DexB^+^ populations. This can be explained by the fact that the BG_D1 sample contained a large population of DexA^+^ cells which showed a different CDR3 region for the TCRα- or TCRβ-chains. Moreover, the motif in Fig. 2E for DexB+ cells is even more distinct than in the TPR-enriched sample, likely due to the presence of a dominant TPR-specific clonotype, as both the TPR-enriched and BG_D2 samples originated from the same donor. This highlights how dextramer labeling effectively reveals distinct CDR3 motifs and dominant clonotypes, especially when the true extent of T-cell heterogeneity is unknown.

Subsequently, we analyzed clonotypes within the dataset by first filtering immune receptor sequences to exclude BCR, VJ or VDJ chain-only sequences, as well as ambiguous and multi-chain sequences. Clonotypes were then defined in two steps. Initially, cells sharing identical CDR3 nucleotide sequences in their primary VJ and VDJ chains were assigned matching clone IDs. Subsequently, a more flexible ‘clonotype cluster’ label was added based on amino acid sequence similarity. This involved calculating a pairwise distance matrix of unique receptor configurations, allowing receptor sequences with CDR3 amino acid distances below a specific cutoff to form a clonotype cluster. This method enabled identification of TCR sequences potentially recognizing the same antigen despite not sharing identical nucleotide sequences.

Clonotypes and clonotype clusters were visualized using network graphs (Figure 3). In these graphs, circles represent clonotypes, and edges between them indicate clonotype clusters. The size of each circle reflects its clone size, and the label specifies the clonotype cluster number. These graphs also integrate information from other data layers.

**Figure 3.**
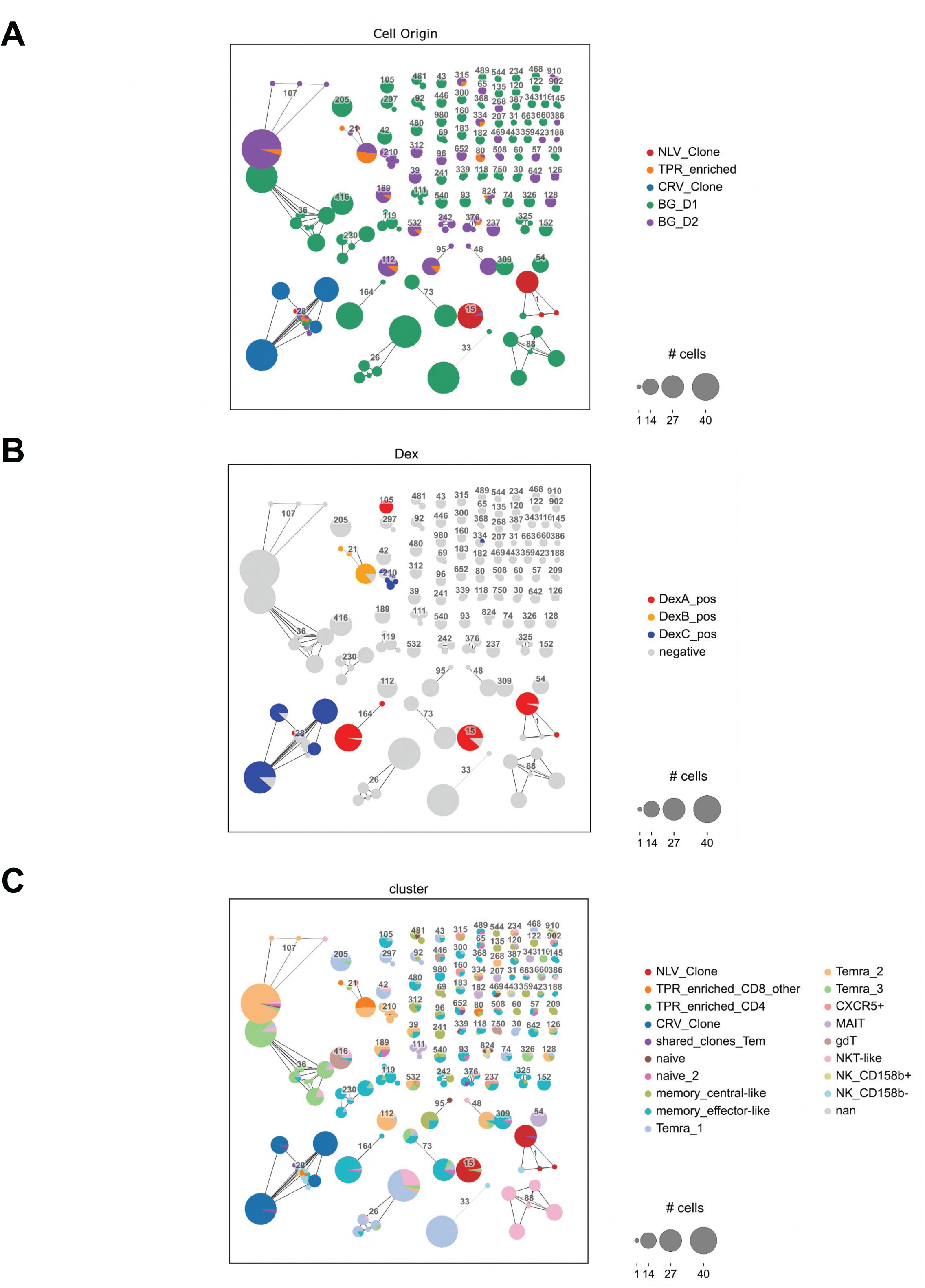
TCR repertoires of CMV-specific CD8^+^ T cells. Clonotype network graphs for filtered TCR sequences are coloured by flex-tag label (A), Dextramer label (B) and annotated cluster from scRNA-seq analysis (C). Each circle depicts a clonotype (defined as identical nucleotide CDR3 sequences of VJ and VDJ chains). The size of circles indicates the clone size. Connecting lines between clones indicate a ‘clonotype cluster’, thus a high degree of CDR3 amino acid similarity between the VJ and VDJ chains of the connected clones. Similarities were based on weighted mismatch distances from protein sequence alignments with a BLOSUM62 substitution matrix. This approach, which is also called TCRdist, was previously developed by Dash et al.^28^. Sequences with a distance metric below 15 are connected by lines (e.g. a distance of 10 is synonymous to 2 R’s mutating into N). The numeric label indicates the clonotype cluster number. Clonotype clusters in the graph layout are arranged according to their size.

Analysis of flex-tag label information showed that cells from the CRV clone were found in clonotype cluster ‘28’, whereas cells from the NLV clone, as predicted from the logo plots of CDR3 sequences, were observed in clusters ‘1’ and ‘15’ (Figure 3A). Cells in the NLV clone sample shared a common TCRβ chain with the CDR3 amino acid sequence ‘CASSPKTGTIYGYTF’, originating from *TRBV6-5, TRBD1*, and *TRBJ1-2* (Table 3). However, their TCRα chains differed: clonotype cluster ‘15’ expressed the CDR3 aa sequence ‘CARNTGNQFYF’ (*TRAV24, TRAJ49*), while clonotype cluster ‘1’ expressed ‘CVRNRDDKIIF’ (*TRAV38-1, TRAJ30*) (Table 3). Cells in the CRV clone shared a common TCRα CDR3 amino acid sequence ‘CAASEAAGNKLTF’ (*TRAV29/DV5, TRAJ17*), with the most prominent TCRβ CDR3 amino acid sequence being ‘CASSPVSQGVRYNEQFF’ (*TRBV28, TRBD1, TRBJ2-1*), followed by ‘CASRCPDRGQGYEQYF’ (*TRBV12-2, TRBD1, TRBJ2-7*) in a smaller portion of cells or as a secondary VDJ chain (Supp. Fig. 4B).

DexA^+^ cells and cells labeled with a flex-tag for the NLV clone co-localized to clonotype clusters ‘1’ and ‘15’ (Figure 3A,B). However, two additional DexA^+^ clusters, ‘105’ and ‘164’, originated from the polyclonal background sample of donor 1 (BG_D1) (Figure 3B). TCR sequences of clusters ‘105’ and ‘164’ differed from those of clusters ‘1’ and ‘15’ of the NLV clone sample (Table 3). Clonotype cluster ‘21’ contained DexB^+^ cells and comprised cells from both the TPR enriched and BG_D2 samples, which was expected since both samples originated from the same donor. Despite the heterogeneity of the TPR enriched sample, DexB staining helped isolate TCR sequences specific for pp65_417-426_ on HLA-B*07:02. The most frequent TCR sequences in DexB^+^ cells were CATVLRMDSSYKLIF (*TRAV17, TRAJ12*) and CASSLLGISTYNEQFF (*TRBV7-9,TRBJ2-1*) (Figure 2E, Supp. Fig. 4B).

Furthermore, the specificities of TCRs identified were assessed by comparing sequences to a curated database of TCR sequences with known antigen specificities (VDJdb). This comparison revealed that the CDR3 sequences of the NLV and CRV clones were correctly predicted to be specific for pp65 and IE-1 CMV epitopes, respectively (Sup. Fig. 4C). In summary, the analysis demonstrated that large clones originated from the three spike-in samples, or, alternatively, from clusters of differentiated T cell populations in the background samples (Figure 3C).

## 6. Discussion

This study presents an exemplary workflow using the BD Rhapsody single-cell platform to collect and analyze paired multi-omics data, including whole transcriptome, 31 surface markers, immune receptor sequences, antigen specificities, and multiplexing labels. This is, to our knowledge, the first reported single-cell multi-omics dataset of this kind using the BD Rhapsody single-cell system, as well as the first which uses a dual hierarchical multiplexing strategy by combining flex tags and standard sample tags, thereby significantly enhancing sample multiplexing capabilities and allowing for the potential scaling-up to screen large sample sizes. The dataset aimed to address both technical and biological questions, such as the efficacy of dCODE dextramers in identifying low-frequency antigen-specific T cells and characterizing novel TCR sequences of human CMV-specific T cells.

The overall dataset included five samples: three enriched for CMV-specific CD8^+^ T cells and two CD8^+^ polyclonal T cell backgrounds from CMV-seropositive individuals with matching HLA allotypes. These samples were labeled with flex sample tags and subsequently counted, pooled and stained with dCODE dextramers and standard sample tags at both low (5%) and high (50%) frequencies of the CMV-enriched samples. Despite lower-than-expected spike-in frequencies, likely due to low cell viabilities after thawing (63% and 80% for the NLV and CRV clone, respectively), 9,771 cells with both sample tag information were identified using the dual multiplexing strategy. Notably, using negative selection for enrichment of CD8 T cells resulted in some non-T cell contamination in the polyclonal background samples.

A particular focus of this study was the detection of antigen-specific T cells using dCODE dextramers as sequencing reagents. One key question was whether dextramers could detect antigen-specific T cells at low frequencies. The results confirmed that dextramers were indeed capable of this, even when spike-in frequencies were below 1%. For instance, the NLV clone in one sample had a frequency as low as 0.4%, yet dextramer-positive cells from this clone were still detectable.

While we found that a detection limit for antigen-specific T cells was not reached at the low spike-in frequencies observed in our data, we saw differences in dextramer sensitivity when used as a sequencing versus flow cytometry reagent. Flow cytometry showed that 97% of NLV clone cells were DexA-PE^+^ and 98% of CRV clone cells were DexC-PE^+^. Cells detected in the sequencing dataset labeled with flex tags for the NLV and CRV clone could therefore be assumed to bind to DexA and DexC, respectively. The sequencing data confirmed the high specificity of DexA and DexC for the NLV and CRV clones, respectively, with negligible false positives and no HLA-mismatched binding. However, the sensitivity of sequencing was lower than flow cytometry, with estimated sensitivities of 81.2% for DexA and 83.1% for DexC. This discrepancy may arise from signal loss during multiple processing steps in the sequencing workflow and needs to be further monitored and investigated in future studies.

Further analysis of TCR sequences from CMV-specific T cell clones revealed two previously described pp65-binding CDR3α sequences: ‘CARNTGNQFYF’ with TCR gene usage *TRAV24* and *TRAJ49* from clonotype cluster 1 (DexA^+^) was reported to bind to pp65_495-503_ on HLA-A*02:01^18^. Moreover, the sequence ‘CATVLRMDSSYKLIF’ (*TRAV17* and *TRAJ12*) from clonotype cluster 22 (DexB^+^) was shown to bind to pp65_417-426_ on HLA-B*07:02 (Dössinger, 2014). Additionally, one sequence from the Cancer Genome Atlas study was identified (‘CAASEAAGNKLTF’ in clonotype cluster 3, DexC^+^), although its antigen specificity was not known. For TCRβ chains, the CDR3β regions ‘CASSLLGISTYNEQFF’ (TRBV7-9, TRBJ2-1) and ‘CASSPVSQGVRYNEQFF’ (TRBV28, TRBJ2-1) have been identified as specific to the CMV epitopes pp65_417-426_ and IE-1_309-317_, respectively^19^. Additionally, the CDR3β sequence ‘CASSPKTGTIYGYTF’ (TRBV6-5, TRBJ1-2) was previously described as specific to a pp65 mini-LCL without epitope determination^19^.

These findings provide valuable insights into planning future studies with dCODE dextramers. We observed that dCODE dextramers can identify antigen-specific CD8^+^ T cells at low frequencies and may be used on samples without prior fluorescence-activated cell sorting. The necessity of enriching antigen-specific T cells prior to sequencing depends on various factors, including the experimental question, available cell numbers, and dextramer characteristics. Flow cytometry can help to estimate antigen-specific T cell frequencies and guide sequencing experiments. In cases where antigen-specific T cells are rare, sorting before sequencing may be advantageous to ensure comprehensive gene expression analysis.

Taken together, this comprehensive single-cell multi-omics approach enhances our understanding of immune responses at the single-cell level, particularly in the context of TCR specificity and function. With this manuscript, we provide a detailed protocol and an analytical workflow for the analysis of such data, which holds significant potential for improving vaccine trials and developing targeted immunotherapies.

## 7. Limitations of the study

Evaluation of the performance of dCODE dextramer is contingent on the methods used to define pMHC multimer specificity. Various approaches exist for identifying dextramer-positive (Dex^+^) cells, each of which has its own merits. However, the lack of standardized best practices in the community hampers the comparability of pMHC multimer-omics data between different studies. Establishing community standards would improve the reliability and reproducibility of these assessments. In our study, we used k-means clustering to identify Dex^+^ cells, which proved effective due to the presence of distinct Dex-high and Dex-low populations. However, this method may encounter challenges in datasets featuring populations with intermediate dexamer affinity or high variability between donors.

The existing literature illustrates the diversity in methodologies for defining pMHC-binding T cells. For instance, the TetTCR-SeqHD method employs a bimodal distribution fitting and knee point analysis to set thresholds for defining tetramer-binding events^20^. Another study classified cells as dextramer-positive if the UMI counts for any dextramer were at least twice as high as those for negative controls^11^. Yet another study defined cells as dextramer-positive if more than 30% of dextramer-derived UMI counts were derived from a specific dextramer barcode, with a threshold of ≤3 UMIs used to exclude cells from epitope specificity assignment^21^. Lastly, ICON, a tool for normalization of high-throughput TCR-pMHC binding data and identification of TCR-pMHC interactions, additionally incorporated negative control dextramers as well as TCRαβ pairing information to determine dexamer-binders^22^. These differences highlight the need for harmonized protocols.

In conclusion, while our study demonstrates the potential of combining dCODE dextramers and single-cell workflows like the BD Rhapsody platform for the identification of antigen-specific T cells, it underscores the necessity for standardized methodologies and highlights the challenges inherent in multi-modal single-cell analysis. Future work should aim to establish best practices and further refine these techniques to enhance their applicability, robustness and reproducibility.

## 8. Methods

### Thawing of human primary cells

Cryovials were thawed at a maximum of two at a time. The vials were removed from liquid nitrogen storage and intermediately stored on dry ice for a short period of time. The thawing process was started by holding vials in water at 37°C for two to three minutes. Thawed cells from a cryovial were gently transferred into a 50 ml falcon tube using a 1 ml pipette tip. The cryovial was rinsed with 1 ml warm complete growth medium (RPMI + 10% FCS). With a 1 ml pipette tip, the rinse solution was slowly added to the falcon drop-wise. Cells were serially diluted with complete growth medium a total of five times by 1:1 volume addition, yielding a final volume of 32 ml cell suspension. Cells were centrifuged at 300 g for five minutes at room temperature. Supernatant was aspirated until 1-2 ml of volume remained. Subsequently the cells were resuspended and stored on ice while viable cells were counted.

### Enrichment of human CD8+ T cells from PBMC samples

For the polyclonal “background” samples used in the multi-omics single-cell workflow, human CD8^+^ T cells were enriched from PBMCs of two different donors. After thawing of PBMC samples and counting, CD8+ T cells were isolated by negative selection using MS columns with a Miltenyi human CD8+ T Cell Isolation Kit (Cat No. 130-096-495) according to manufacturer’s instructions. Sterile filtered PBS with 0.5% BSA and 2 mM EDTA was used as a buffer for magnetic cell separation.

### Multi-omics staining and single-cell capture protocol

A total of three staining steps (A-C) were performed prior to single-cell capture and lysis with the BD Rhapsody cartridge (Suppl. Figure 1). Samples were thawed and counted as previously described. Three samples contained CD8+ T cell clones or cells stimulated and expanded by peptide stimulation. Subsequently, they will be called ‘spike-in samples’. Furthermore, CD8+ T cells were isolated from the two PBMC samples as previously described and counted. In the following these are called ‘Background Donor 1’ (HLA-A*02:01+ & HLA-B*07:02/C*07:02-) and ‘Background Donor 2’ (HLA-A*02:01- & HLA-B*07:02/C*07:02+).

#### First sample multiplexing and CD158b staining (Part A)

All five samples were individually labeled in the first step using flex sample tags. Cell suspensions with three to four million cells per sample were transferred to 5 ml LoBind Eppendorf tubes and spun down for 5 minutes at 400g. Supernatants were removed and cell pellets were resuspended in 100 μl of blocking buffer (95 μl BD Stain Buffer (FBS) and 5 μl FcR human blocking reagent) and incubated on ice for 10 minutes. Next, 7.5 μl of primary antibody (CD45-PE; BioLegend, Cat No. 304008) was added on top and the mixture was incubated on ice for 20 minutes. After the incubation, cells were washed three times by adding 2 ml of stain buffer and centrifuging at 400 g for 5 minutes. After the last wash, supernatant was discarded, and the cells were resuspended in 80 μl of stain buffer and placed on ice.

Next, all five samples were labeled with secondary antibodies, namely flex sample tags, which bind to phycoerythrin (PE). For each sample, a sample tag with a different oligo barcode was used. In the same staining, cells were also labeled with an CD158b AbSeq antibody (BD, Cat. No. 559784). 98 μl of stain buffer, 20 μl of flex sample tag and 2 μl of CD158b antibody were added to each sample and incubated on ice for 30 minutes. Next, cells were washed three times by addition of 2 ml of stain buffer and centrifugation at 400 g for 5 minutes with subsequent removal of the supernatant. Finally, cell pellets were resuspended in 500 μl of stain buffer and stored on ice.

#### dCODE Dextramer and second sample multiplexing staining (Part B)

Before the start of the Dextramer staining, the following solutions were prepared: 1) Cell labeling buffer made of BD Stain Buffer (FBS) (BD, Cat No. 554656) with 0.1 g/l sheared herring sperm DNA (Invitrogen, Cat No. 15634017), and 2) Wash buffer made of PBS (pH 7.4) with 2.5 % FBS.

Cells were counted and pooled into four samples. All samples contained a total of 400,000 cells in 50 μl cell labeling buffer and were transferred to 5 ml LoBind Eppendorf tubes. Each sample consisted of either 95 % or 50 % background and 5 % or 50 % spike-in, respectively. For pooling, either ‘Background Donor 1’ or ‘Background Donor 2’ was used as the background. The spike-in always consisted of all spike-in samples (NLV Clone, Enriched TPR and CRV Clone) in a 1:1:1 ratio.

A dextramer labeling master mix was prepared for all four samples consisting of 3.2 μl of 100 μM d-Biotin (diluted in PBS) and 8 μl of each dextramer. The exception was the HLA-C*07:02 (CMV IE-1) dextramer, which was added at 40 μl. Then, 16.8 μl of master mix was added to each sample. Cells were incubated at room temperature for 10 min. Afterwards, 113.2 μl of stain buffer and 20 μl of standard sample tag were added to each reaction. A different sample tag was used for each sample. The mixtures were incubated at room temperature for 20 minutes. Samples were washed three times by addition of 2 ml of stain buffer, centrifugation at 400 g for 5 minutes and removal of supernatant. Lastly, cell pellets were resuspended in 500 μl of stain buffer, counted and stored on ice.

#### AbSeq surface marker staining (Part C)

Before starting, BD AbSeq Immune Discovery Panel (IDP) antibodies (BD, Cat No. 625970) were brought to room temperature for 5 minutes and then spun down. 35 μl of nuclease-free water was added to the bottom of the tube and the antibody mix was reconstituted for another 5 minutes at room temperature, then stored on ice intermediately.

Next, 100,000 cells of each sample were pooled 1:1:1:1 in a 5 ml LoBind Eppendorf tube. Samples were centrifuged at 300g for 5 min at 4°C. Supernatants were discarded and cells were resuspended in 100 μl of blocking buffer. 65 μl of stain buffer was added to the antibody solution for a total volume of 100 μl 2x AbSeq labeling master mix. 100 μl of master mix was added to the cells and the mixture was incubated for 40 min on ice. Afterwards, the sample was washed twice with 2 ml of BD stain buffer as described before. Next, cells were resuspended in 300 μl of cold sample buffer. Finally, 20,000 cells were processed on a BD Rhapsody cartridge with enhanced cell capture beads.

### Multi-omics library preparation

From each cartridge, six libraries were prepared. Reverse transcription and template switching was performed following manufacturer’s instructions (Becton Dickinson, Doc ID: 23-24020(02)). During the subsequent denaturation, 75 μl of elution buffer was used. The denatured supernatant was saved and utilized for dCODE Dextramer (RiO), AbSeq and Sample Tag library production. Generation of these three libraries was done following the manufacturer’s protocol (Immudex, Doc ID: TF1196.07) with 11 instead of 13 cycles in the dCODE RiO PCR2. By contrast, indexed WTA, TCR and BCR sequencing libraries were generated directly from the cell capture beads by following the corresponding steps of the manufacturer’s protocol (Becton Dickinson, Doc ID: 23-24020(02)).

### Sequencing and raw data demultiplexing

Libraries were pooled taking into account the lower clustering efficiency of BCR and TCR libraries due to the increased average length following manufacturer’s instructions (Becton Dickinson, Doc ID: 23-24020(02)). Sequencing was performed on a NovaSeq6000 instrument with a S4 300 cycles kit v1.5. Sequencing was performed paired end with read 1 or 86 bp and read 2 216 bp for optimal TCR/BCR assembly. Raw sequencing output was demultiplexed and converted to FASTQ files with bcl2fastq2 v.2.20.

### Analysis of paired data for whole transcriptome, surface markers, sample tags and dCODE dextramers

Preprocessing of the raw sequencing data of everything except AIRR libraries was performed with a pipeline, which was based on the Drop-seq tools from McCarroll Lab (version 0.4, available at https://github.com/Hoohm/dropSeqPipe). These were originally developed for the processing of Drop-seq data^23^ but were adapted to work on BD Rhapsody’s enhanced cell capture bead structure. WTA libraries were STAR aligned to the human GENCODE reference genome and transcriptome hg38 release 33^24^. Sample tag, AbSeq and RiO dCODE Dextramer libraries were aligned to their respective barcode reference sequences.

The resulting UMI count matrices were imported into R (version 4.1.2) as previously described^25^ and further analyzed with the R toolkit Seurat (version 4.1.0)^16^. RNA-seq analysis using Seurat was performed with the docker image jsschrepping/r_docker:jss_R412_S41 (https://hub.docker.com/r/jsschrepping/r_docker). The WTA count matrix was filtered to only contain cells with more than 250 UMI counts and with a percentage of mitochondrial genes of less than 55 %. Sample tag count matrices were split into separate assays for flex and regular sample tags. Dextramer count matrices were obtained for four dextramers – three HCMV-specific ones (DexA, DexB and DexC) and one negative control dextramer (DexN). Background noise correction was performed by subtracting counts of the DexN Dextramer from DexA, DexB and DexC counts on a per-cell basis.

Count matrices for whole transcriptome, AbSeq markers, flex sample tags, regular sample tags and background corrected Dextramers were merged into one Seurat object with distinct assays using the CreateAssayObject function. The count matrix of the whole transcriptome assay was normalized using sctransform v2 regularization (R release 0.3.3) (Hafemeister and Satija, 2019). The count data in the assays for AbSeq, sample tags and background-corrected Dextramers were normalized using centred log-ratio (CLR) transformation followed by scaling. For WTA and AbSeq assays, principal components (PCs) were calculated with Seurat’s RunPCA function.

Demultiplexing of cells was performed for both the flex and regular sample tag assay using the HTODemux function implemented in Seurat^26^. Only cells that were labelled as “singlets” during demultiplexing were kept. Cells were further annotated according to their sample tag labels (Table 1).

Seurat’s FindMultiModalNeighbors function was used to create an embedding that represented a weighted combination of the two modalities ‘whole transcriptome’ and ‘AbSeq’^16^. The first 20 dimensions of the WTA PCA and the first 18 dimensions from the AbSeq PCA were used to calculate multimodal neighbours as well as a weighted shared nearest neighbour (WSNN) graph. The resulting multimodal neighbour object was used to compute a joint weighted-nearest neighbours Uniform Manifold Approximation and Projection (wnnUMAP) embedding with Seurat’s RunUMAP function. The WSNN graph was used to identify clusters with the leiden algorithm (resolution 0.5). Gene and surface expression markers for each cluster were determined with a Wilcoxon rank sum test using the functions FindAllMarkers function (with the following parameters: only.pos = TRUE, min.pct = 0.25, logfc.threshold = 0.25). In case of the sctransform-normalized WTA assay, the function PrepSCTFindMarkers was applied to the Seurat object prior to marker identification. Data visualization was performed with Seurat’s VlnPlot, DotPlot, DoHeatmap, FeaturePlot and DimPlot function, as well as with R package ggplot2 (version 3.3.5). Dextramer-positive cells were defined using k-means clustering applied to CLR-normalized and background corrected counts for each Dextramer individually. This was done using the kmeans function (centers = 4) from the stats package (version 4.1.2). The lower boundary of the k-means cluster with the highest mean value was used as a threshold to separate dextramer-positive from dextramer-negative cells. Dextramer sensitivities and specificities for the DexA & the NLV clone, as well as DexC & the CRV clone, were calculated with the confusionMatrix function of the R package caret (version 6.0-94).

### Analysis of paired TCR/BCR data

The raw fastq files of the TCR and BCR libraries were preprocessed with MiXCR (version 4.2.0) using the mixcr analyze function with the ‘bd-rhapsody-human-tcr-full-length’ or ‘bd-rhapsody-human-bcr-full-length’ preset, respectively. AIRR compliant tables were exported with MiXCR’s exportAirr function.

The subsequent downstream analysis of the paired TCR/BCR and WTA data was performed in Python (version 3.10.13) with the docker image rnakato/shortcake_light_3.0.0 (https://hub.docker.com/r/rnakato/shortcake_light). The Scirpy toolkit (version 0.17.2) was used for the subsequent analysis and visualization of scAIRR-seq data as well as the integration with the corresponding scRNA-seq data^27^. Additionally, the following python packages were used: Scanpy (1.9.8), anndata (0.10.8), muon (0.1.6), mudata (0.2.3), numpy (1.26.4), scipy (1.13.0), pandas (2.2.2) and matplotlib (3.7.2).

The AIRR-compliant output tables were imported into Scirpy with the scipry.io.read_airr function. Chain quality metrics were calculated with the function ir.tl.chain_qc. In order to use the filtered, normalized and annotated Seurat object for downstream analysis with Scirpy and Scanpy, it was converted into an AnnData object with the SeuratDisk R package (version 0.0.0.9019). It was imported with the scanpy.read_h5ad function. Transcriptome data was merged with TCR annotations by creation of a multimodal data object with the MuData command. For subsequent clonotype construction, the immune receptor repertoire was filtered to exclude sequences that fell into the chain pairing categories “ambiguous”, “multichain”, “no IR”, “orphan VJ” and “orphan VDJ”, as well as any entries that contained BCR sequence annotation.

Clonotypes were defined based on CDR3 nucleotide sequence identity of primary VJ and VDJ chains with the functions scirpy.pp.ir_dist and scirpy.tl.define_clonotypes. Clonotype networks were defined on CDR3 amino acid sequence identity using the functions scirpy.pp.ir_dist(metric=”alignment”, sequence=”aa”, cutoff=15) and scirpy.tl.define_clonotype_clusters (sequence=”aa”, metric=”alignment”, receptor_arms=”all”, dual_ir=”any”). TCR CDR3 sequence logo plots were created with palmotif (version 0.4). Specificity prediction analysis was done by VDJdb database query using the function scirpy.tl.ir_query with parameters: metric=”identity”, sequence=”aa”, receptor_arms=”any”, dual_ir=”any”. The multimodal data object was annotated using scirpy.tl.ir_query_annotate with strategy “most-frequent”.

### Data availability

The accession numbers for the data generated in this paper are as follows: single-cell RNA sequencing data can be accessed under EGAXXX.

## 9. Acknowledgements

We thank the PRECISE team for technical assistance and Tal Pecht and Vadir Lopez-Salmeron (BD Biosciences) for critical discussions and technical support.

Anna C. Aschenbrenner is a member of the excellence cluster ImmunoSensation2 (EXC 2151) funded by the German Research Foundation (DFG) under grant agreement #390873048) and is supported by the DFG via the SFB 1454 – project number #432325352; grant #458854699; grant #466168337; grant #466168626; the BMBF-funded project IMMME (01EJ2204D); and the EU-funded project ImmunoSep (#847422) and NEUROCOV receiving funding from the RIA HORIZON Research and Innovation under GA No. 101057775. Joachim L. Schultze is supported by the excellence cluster ImmunoSensation2 (EXC 2151); the EU-funded projects discovAIR (#874656) and SYSCID (#733100) and NEUROCOV receiving funding from the RIA HORIZON Research and Innovation under GA No. 101057775; the BMBF-founded project Diet-Body-Brain (DietBB, 01EA1809A); the DFG via the SFB 1454 (#432325352) and iTREAT (01ZX1902B). Marc Beyer is supported by the excellence cluster ImmunoSensation2 (EXC 2151, #390873048); the DFG via the IRTG2168 (#272482170), SFB1454 (#432325352); the EU-funded project NEUROCOV receiving funding from the RIA HORIZON Research and Innovation under GA No. 101057775; the Else-Kröner-Fresenius Foundation (2018_A158). Lorenzo Bonaguro is supported by the excellence cluster ImmunoSensation2 (EXC 2151, #390873048) and the DFG-funded project ImmuDiet (#513977171).

## 10. Author contributions

Conceptualization, L.B., A.C.A., J.L.S., A.M., M.D.B.; Data acquisition, analysis, and interpretation, I.G., L.B., M.B., S.M., C.J., E.D.E.; Resources and discussion, A.C.A., A.M.; Supervision and conceptual discussion, L.B., J.L.S., M.D.B., Writing - original draft, I.G., L.B., M.D.B.; Editing, all authors.

## 11. Declaration of interest

The authors declare that there are no competing interests.

## Notes

### Competing Interest Statement

The authors have declared no competing interest.

