## Supplementary Figures for "Characterizing human CMV-specific CD8^+^ T cells using multi-layer single-cell omics"

### 13. Supplementary Figure Legends

**Figure S1. Single-cell multi-omics workflow allows delineation of spike-in and background samples, related to Figure 1.**

Overview of stainings in experimental multi-omics workflow. In total, three staining steps were performed before single-cell capture. (A) First, clones and CD8 polyclonal backgrounds were labeled with flex sample tags and a custom CD158b antibody. (B) Next, clones and CD8 polyclonal backgrounds were counted and pooled in indicated ratios. Staining mixes consist of different combinations of CMV spike-in mix (1:1:1 ratio of NLV & CRV clone, as well as TPR enriched cells) and $CD8^+$ polyclonal T cells from two background donors, either BG_D1 or BG_D2. These samples were stained with dextramers and standard sample tags. All four samples were stained with DexA, DexB, DexC and DexN. (C) Lastly, all four samples were pooled and stained with the BD AbSeq IDP panel before proceeding with single-cell capture.

**Figure S2. Single-cell multi-omics workflow allows delineation of spike-in and background samples, related to Figure 1.**

(A-B) Superimposition of flow cytometry data of individual single-Dextramer stainings (DexA, DexB, DexC, DexN and negative control without Dextramer) on three CMV enriched input populations (NLV clone, TPR enriched and CRV clone). (A) Gating strategy for the quantification of live, single, CD3+ and Dextramer-PE+ cells. (B) Gating for the identification of Dextramer-PE+ cells for four Dextramers (DexA, DexB, DexC and DexN) in three different samples with enriched CMV-specific cells (NLV Clone, Enriched TPR and CRV Clone).

(C-D) Plots originating from single-cell omics data. (C) wnnUMAP representation with cells colored by cartridge origin (technical replicates). (D) Dotplot displaying expression of the top 5 marker genes for each cluster. The dots are coloured by z-transformed mean expression per cluster. The dot size represents the frequency of expression within the cluster.

**Figure S3. Characterization of CMV-specific $CD8^+$ T cells, related to Figure 2.**

(A) Histogram and density plots for distribution of noise-corrected CLR-normalized DexA, DexB and DexC counts. Blue line indicates cutoff for $Dex^+$ label (3.81 for DexA, 3.79 for DexB and 3.87 for DexC), as determined by k-means clustering (k=4). The cutoff was defined as the lowest value of the cluster with the highest mean expression.

(B) Dextramer-positive cells in each staining condition. Barplots indicating frequency of $Dex^+$ cells (for DexA, DexB and DexC) in each of the four staining conditions: Spike-In to Background Ratio of 05:95 and 50:50, with background originating from Donor 1 (BG_D1, HLA-matched to DexA) or Donor 2 (BG_D2, HLA-matched to DexB and DexC).
(C) Pie charts showing cluster origin of $Dex^+$ cells.
(D) Co-expression of DexC and CD158b (AbSeq) per cell. CLR-normalized expression values are plotted and cells defined as $DexC^+$ are highlighted.

**Figure S4. Characterization and TCR repertoires of CMV-specific $CD8^+$ T cells, related to Figure 2 and 3.**
(A) Immune receptor sequence types in the dataset. Barplot depicting cell numbers categorized by type of chain pairing. Applies only to cells, which had both a recorded immune receptor sequence and paired RNA-seq information.
(B) CDR3 sequence logos depicting under- and over-represented amino acids of CDR3α and CDR3β amino acid sequences. Logo plots are shown for samples carrying the flex-tag labels for the NLV clone, TPR enriched and CRV clone input material. Colours indicate amino acid chemistry.
(C) Antigen specificity predictions for the NLV and CRV clones, as well as $DexB^+$ cells from the TPR-enriched sample, were made by matching CDR3 aa VJ and VDJ sequences to the VDJdb reference database. A match was identified if similarity was found in either the VJ or VDJ chain.

**Figure S5. Multimodal single-cell analysis workflow integrating paired and dual-multiplexed RNA, AbSeq, Dextramer, and VDJ sequencing data, related to Methods.**

The workflow outlines the analysis pipeline for paired RNA, AbSeq, Dextramer, Multiplex, and VDJ (primarily TCR) single-cell omics data. With the exception of VDJ data, which was preprocessed using MiXCR, all other data types were processed using a modified Drop-seq pipeline with downstream analysis in Seurat (R).
For whole transcriptome ("RNA") data, only cells with more than 250 unique molecular identifiers (UMIs) and less than 55% mitochondrial gene content were retained. Gene expression was normalized using SCTransform v2, followed by principal component analysis (PCA). Data derived from oligo-conjugated antibodies ("AbSeq") for cell surface proteins was normalized using centered log-ratio (CLR) transformation, also followed by PCA. The RNA and AbSeq data were integrated using a weighted nearest neighbor (WNN) approach, generating a combined UMAP

embedding using the top 20 principle components (PCs) from the RNA and the top 18 PCs from the AbSeq data. Leiden clustering was performed at a resolution of 0.5, and cluster markers for gene expression and surface proteins were identified using the Wilcoxon rank-sum test.
For Dextramer data, background correction was applied by subtracting negative control dextramer (DexN) counts from HCMV-specific dextramer counts. The corrected data was CLR-normalized, and thresholds for Dextramer-positive cells were identified using k-means clustering.
For sample demultiplexing ("Multiplex info"), flex and sample tag counts were separated into distinct Seurat assays, CLR-normalized, and demultiplexed using Seurat's HTODemux function. Only cells, which were categorized as "Singlet" in both assays were kept for further analysis.
VDJ data and the h5ad-converted multimodal data from the Seurat workflow were analyzed using the scirpy toolkit (Python). Cells with matching barcodes in the Seurat object were retained for downstream VDJ analysis (7,499 cells). For chain pairing quality control (QC), cells with ambiguous chains, such as TRA+TRD or TRA+IGH combinations, as well as cells with more than two VJ or VDJ chains ("multichain"), no immune receptor chains ("no IR"), or single-chain cells ("orphan VJ/VDJ") were removed. Clonotypes were defined by CDR3 nucleotide identity, and clonotype networks were built based on CDR3 amino acid sequence similarity. Antigen specificity predictions were made by comparing CDR3 motifs to the VDJdb epitope database.

### Suppl. Figure 1

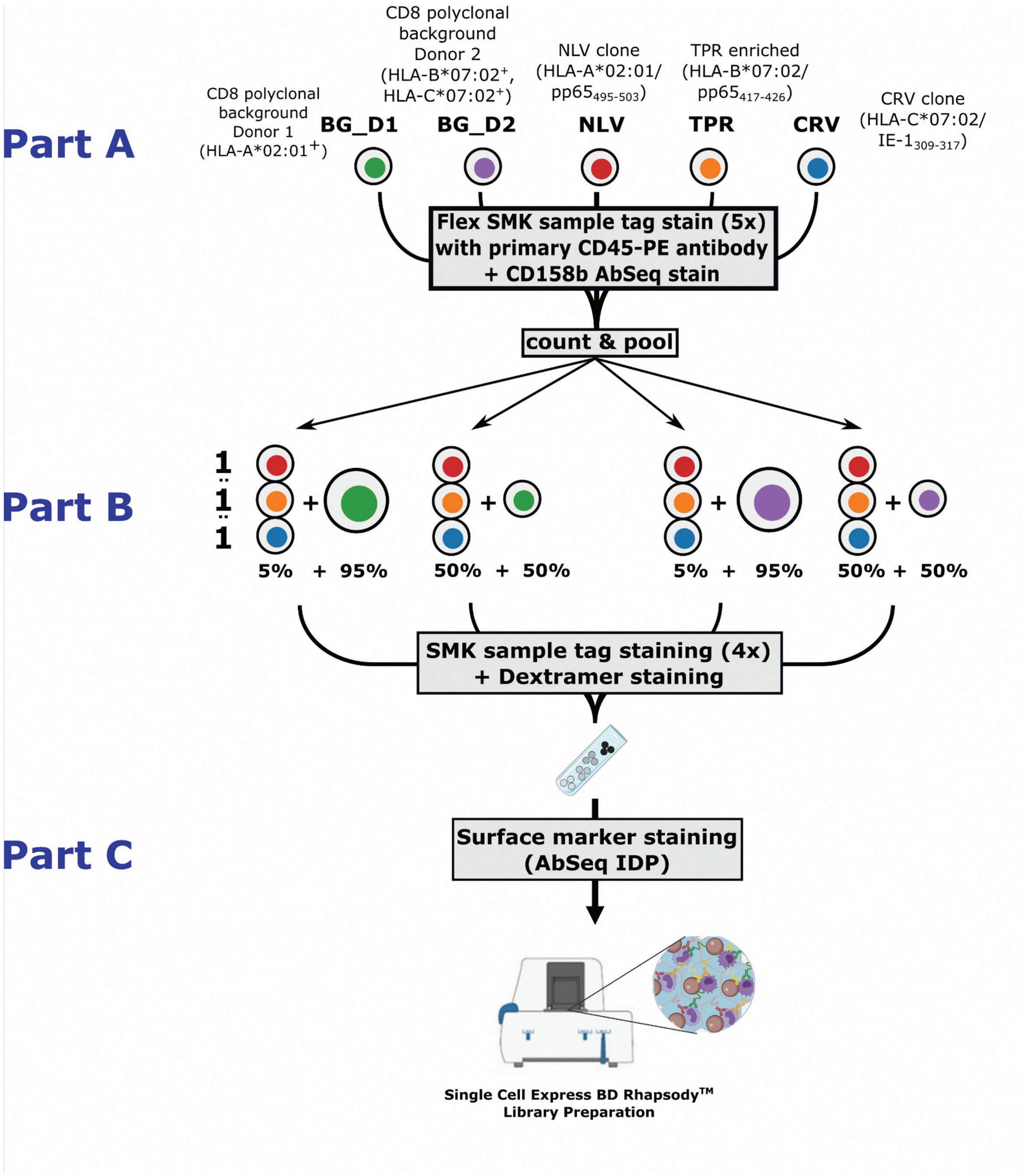

### Suppl. Figure 2

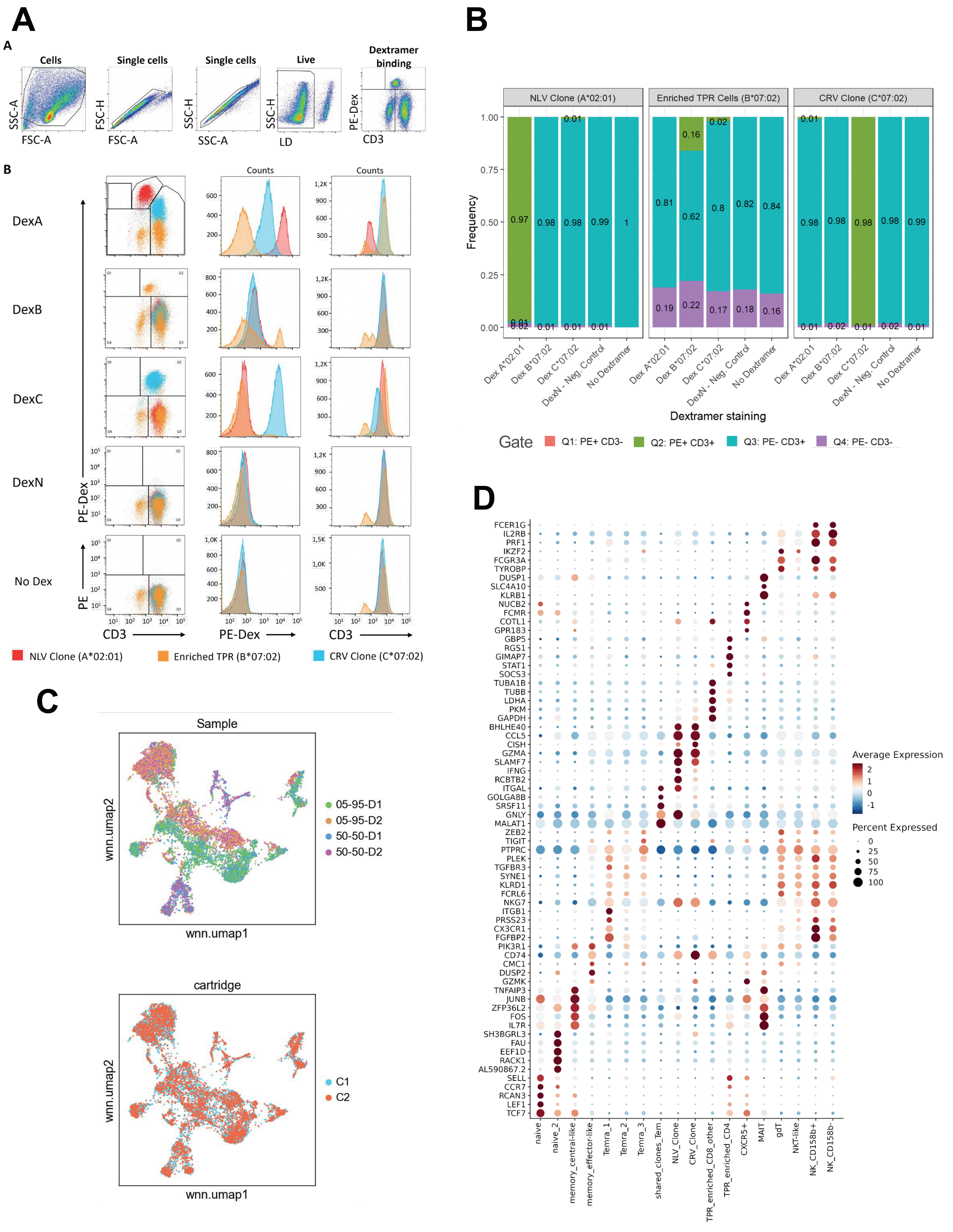

### Suppl. Figure 3

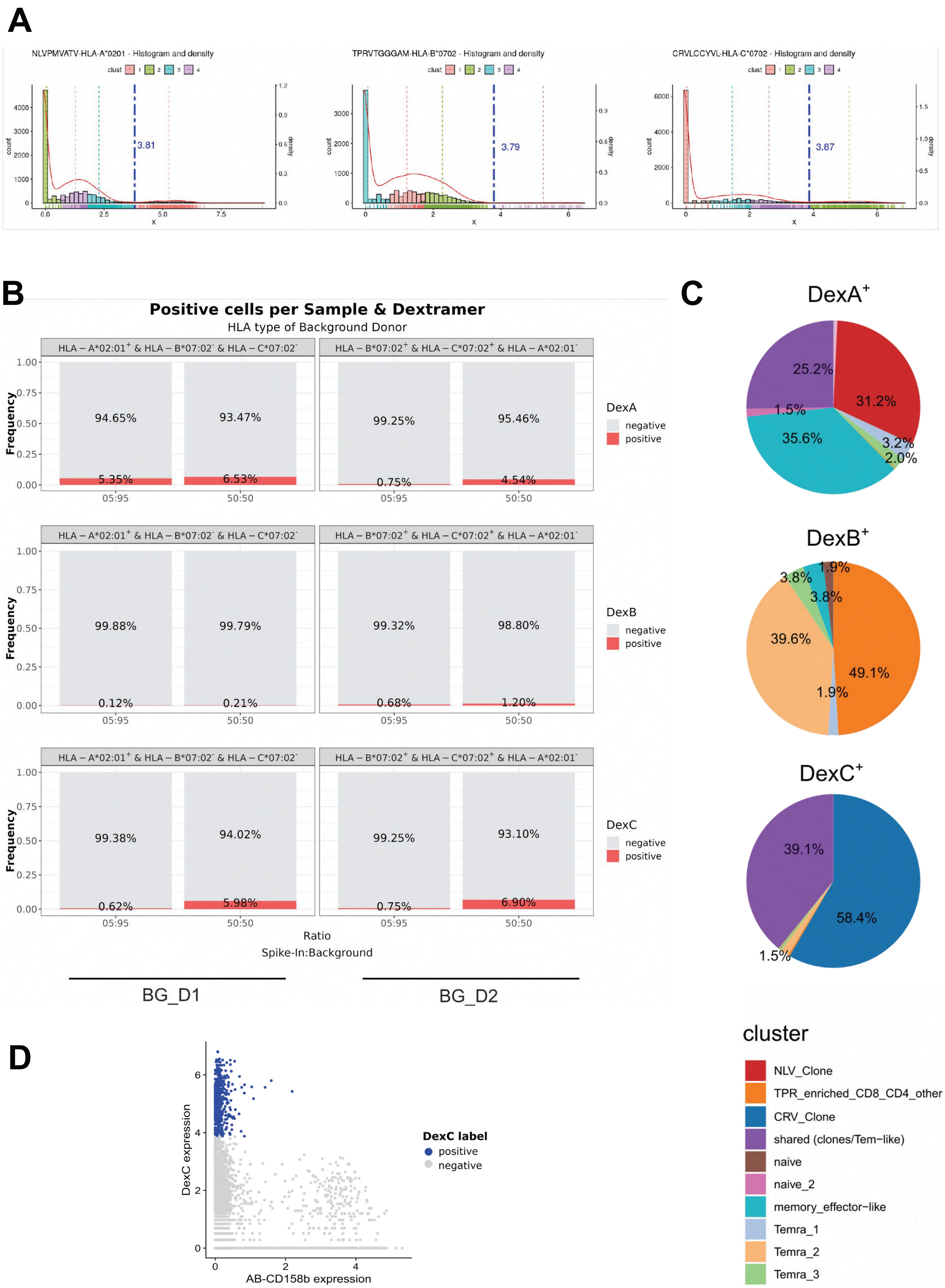

### Suppl. Figure 4

A

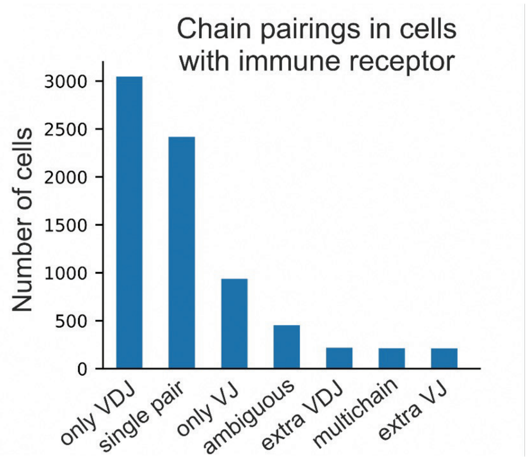


B

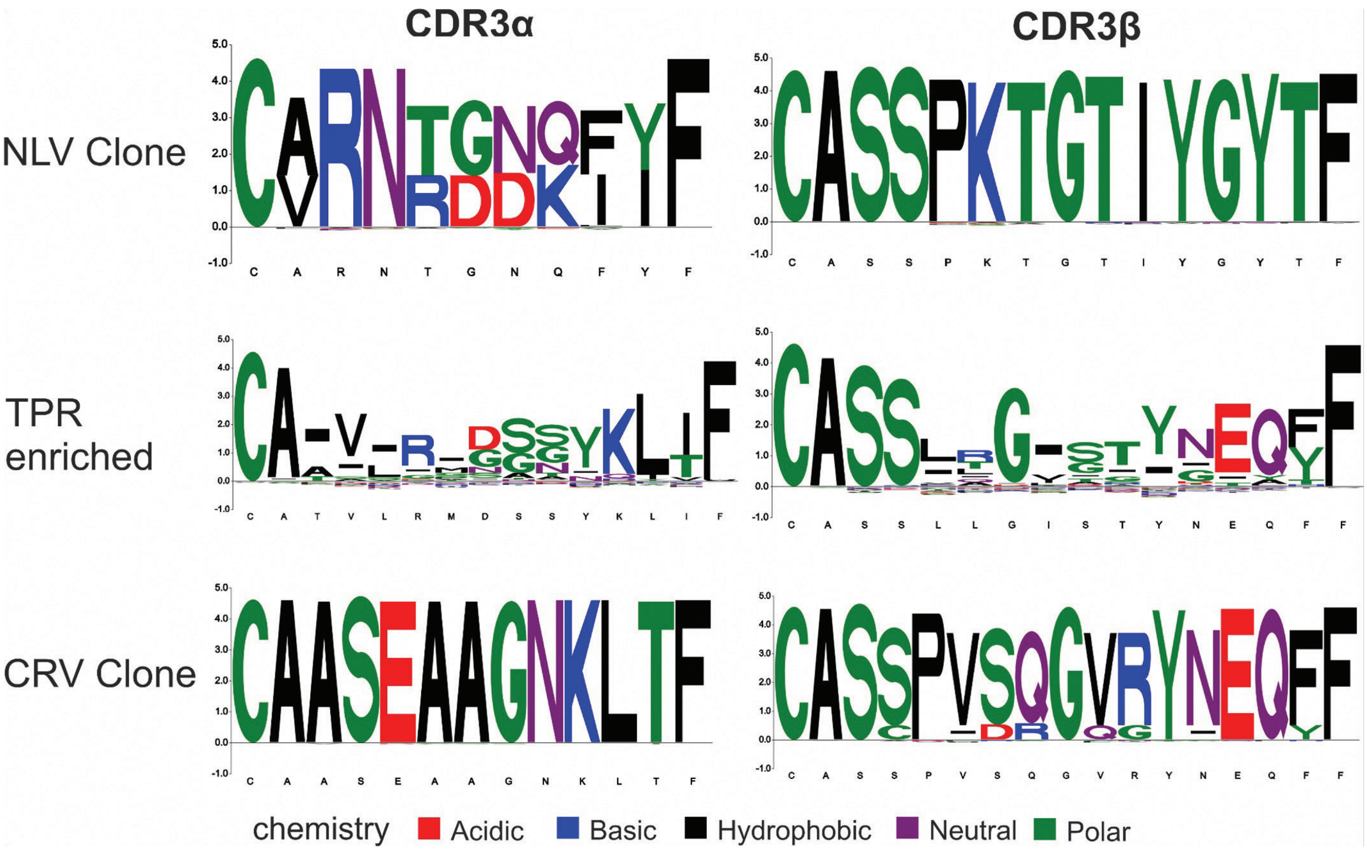


C

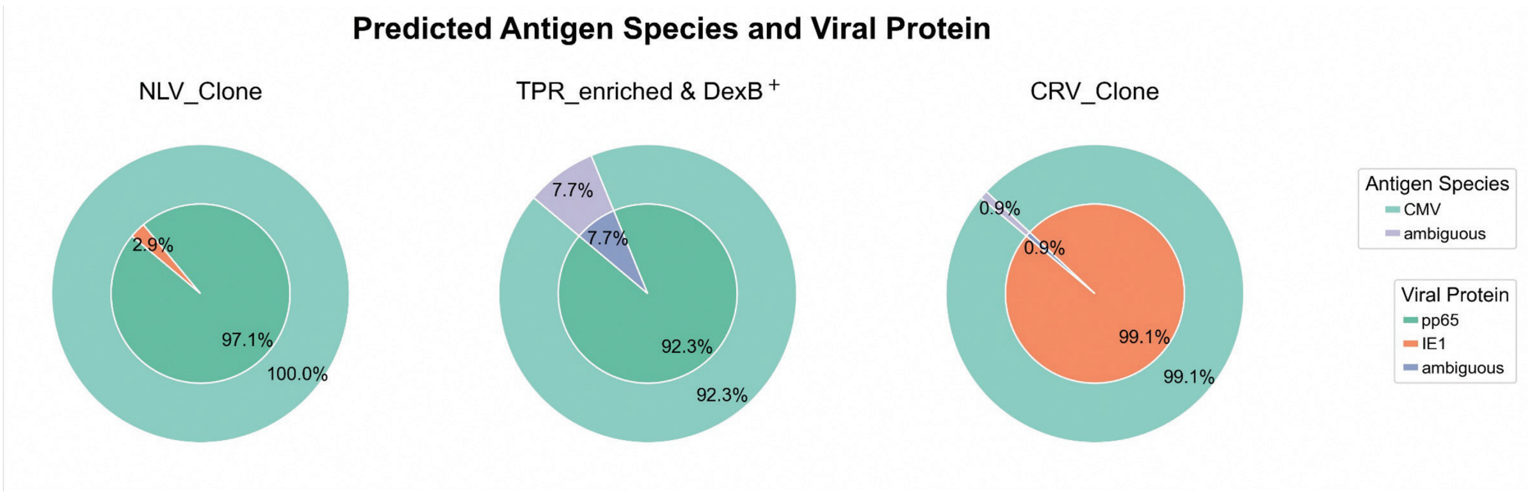

### Suppl. Figure 5

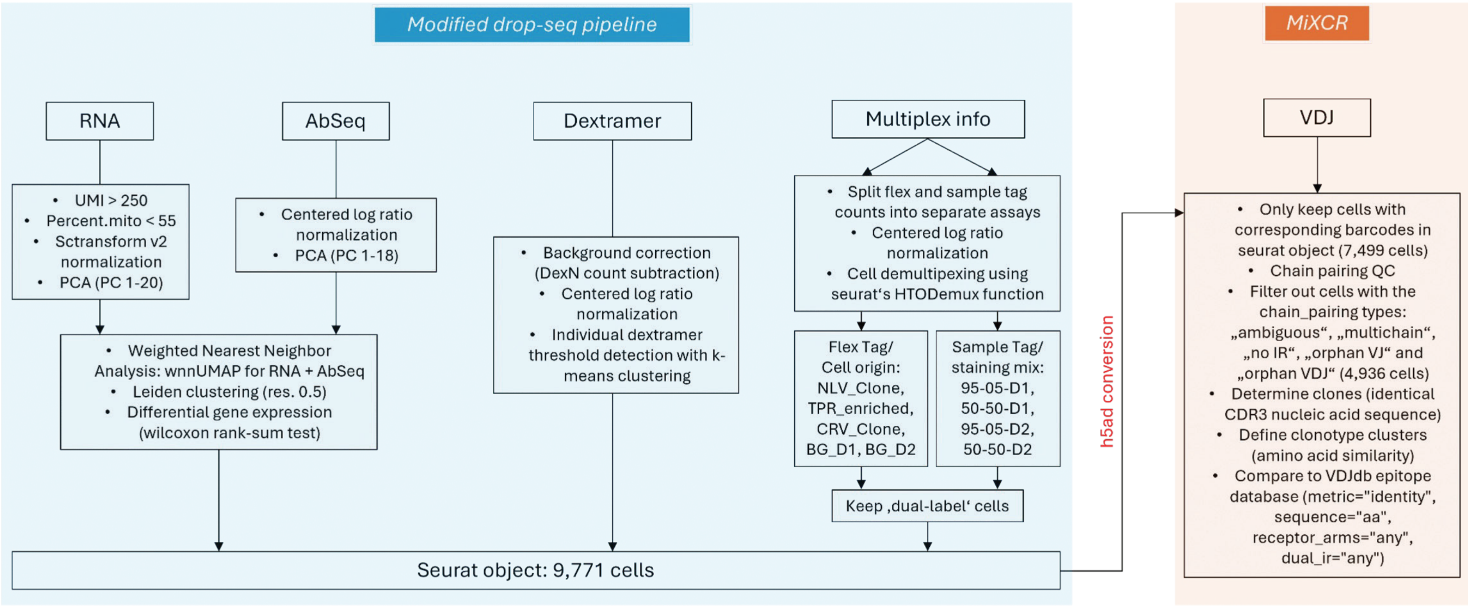
